# Spinal cord injury in rats disrupts thermoregulation and suppresses stress-induced hyperthermia

**DOI:** 10.1101/2025.03.03.641238

**Authors:** John C. Aldrich, Kalina J. Dusenbery, Linda R. Watkins, Andrew D. Gaudet

**Affiliations:** Department of Psychology, College of Liberal Arts, University of Texas at Austin, Austin, TX 78712, USA; Department of Neurology, Dell Medical School, University of Texas at Austin, Austin, TX 78712, USA; Department of Psychology and Neuroscience, and the Center for Neuroscience, University of Colorado, Boulder, CO 80301, USA; Interdisciplinary Neuroscience Program, University of Texas at Austin, Austin, TX 78712, USA

**Keywords:** spinal cord injury, neurotrauma, thermoregulation, autonomic dysfunction

## Abstract

Spinal cord injury (SCI) in humans can robustly dysregulate healthy autonomic nervous system function, including thermoregulation. Despite this, the relationship between SCI and autonomic dysfunction remains incompletely understood and is often overlooked in rodent models that emphasize locomotor recovery. One notable autonomic output in rodents is stress-induced hyperthermia, which is a transient core temperature increase caused by an acute stressor. Here, we tested whether SCI in rats dysregulates stress-induced hyperthermia. To assess SCI-induced thermoregulatory dysfunction, we continuously monitored core temperature in male and female rats using implantable telemetry devices before and after a T8 contusion SCI (or sham surgery including laminectomy). Prior to surgery, stress—in response to handling and cage changes— resulted in transient hyperthermia that peaked ∼1°C higher than baseline and lasted 60-90 minutes. Sham-operated rats retained typical stress-induced hyperthermia beginning immediately after surgery. In contrast, SCI transiently abolished stress-induced hyperthermia in both sexes, indicating a profound disruption in autonomic regulation acutely after injury. Within 3-4 weeks after SCI, the stress-induced hyperthermic response gradually returned and reached pre-injury levels by week seven. Therefore, thoracic SCI in rats abolishes stress-induced hyperthermia, which gradually recovers over time post-injury. Overall, this study underscores the impact of incomplete SCI on autonomic function and highlights the need for future research focused on autonomic outcomes.

**Highlights:**

- Handling and care procedures elicit stress-induced hyperthermia in rats
- Stress-induced hyperthermia was abolished in rats after spinal cord injury
- Stress-induced hyperthermia gradually returned over the subsequent weeks
- By 7 weeks, the hyperthermic response had returned to pre-injury levels

## 1. Introduction

The autonomic nervous system (ANS) regulates the function of nearly all tissues by facilitating communication of sensorimotor information between supraspinal and peripheral regions (Wehrwein et al., 2016, Roche et al., 2023). Through this crosstalk, the ANS governs essential physiologic processes, such as heart rate (Fisher et al., 2015, Sapru, 1991), blood pressure (Lanfranchi & Somers, 2002; Gross et al., 2005; Fu et al., 2006), and digestion (Rogers et al., 1996), thereby ensuring the body maintains internal optimal function (*homeostasis*) in response to external changes. A key function of the ANS is thermoregulation–a process in which the ANS detects temperature deviations and activates involuntary physiologic mechanisms such as blood vessel dilation, sweat gland activity, and metabolic heat production to sustain a stable core body temperature (Morrison & Nakamura, 2011).

Spinal cord injury (SCI) disrupts communication between supraspinal centers and peripheral tissues, impairing myriad physiologic systems, including the ANS (Henke et al., 2022; Alizadeh et al., 2019; Krassioukov, 2009; Wecht et al., 2021). Indeed, over 50% of individuals with high-level SCI experience autonomic dysreflexia, a condition characterized by exaggerated, involuntary increases in blood pressure in response to noxious stimuli below the level of injury (Karlsson, 1999; Krassioukov et al., 2021; Mironets et al., 2018). Similarly, SCI disrupts homeostatic thermoregulatory mechanisms due to disrupted afferent input to the brain’s thermoregulatory center and reduced mediation of effector tissues below the injury site (Downey et al., 1967; Laird et al., 2006; Price and Campbell, 2003). The loss of ANS regulation increases the risk of hyper– and hypothermia in individuals with SCI (Guttmann et al., 1958; Price & Campbell, 1999; Price & Campbell 2003).

Although it is clear that SCI disrupts homeostatic thermoregulation, how SCI affects allostatic processes in thermoregulation remains less understood. Unlike homeostasis, which maintains a fixed physiologic set point, allostasis allows for dynamic adjustments in response to external stressors or demands (Ramsay & Woods, 2014).

One example of an allostatic response is fever, when the body raises its thermal set point to enhance immune function during infection (Saper & Breder, 1994). Similarly, stress-induced hyperthermia occurs when acute psychological or physiological stress elicits a transient increase in core body temperature (Lkhagvasuren et al., 2011; Nakamura & Morrison, 2022; Oka, 2015). These flexible, context-dependent responses illustrate how thermoregulation extends beyond homeostatic reflexes. Thus, there is a need to understand the extent that SCI affects allostatic thermoregulatory processes, such as stress-induced hyperthermia.

Here, we explore whether SCI in rats disrupts stress-induced hyperthermia. Core body temperature was monitored in female and male rats prior to and after thoracic SCI (or sham surgery) (Gaudet et al., 2018). In rats prior to surgery or with sham surgery, care-related handling and other interactions caused pronounced temperature spikes immediately following room entry. SCI (but not sham surgery) abolished these stress-induced hyperthermic responses, suggesting widespread autonomic and thermoregulatory dysfunction. SCI rats gradually recovered stress-induced hyperthermic responses over 3-4 weeks. Thus, clinically relevant SCI in rats perturbs typical stress-related thermoregulation, particularly acutely after injury.

## 2. Materials and methods

### 2.1. Surgery, post-operative care, and temperature recording

These experiments were approved by the University of Colorado Boulder Institutional Animal Care and Use Committee and conformed with ARRIVE (Animal Research: Reporting In Vivo Experiments) standards. The data analyzed in this study were collected as part of a larger study (Gaudet et al., 2018). Male and female Sprague Dawley rats (2–3 months old; Evigo) were individually housed on a 12-hour light/dark cycle with ad libitum food and water. Rats were implanted intraperitoneally with radiotelemetric transmitters (MiniMitter, Respironics) two weeks before surgery to continuously monitor core body temperature using TR-4000 receiver boards and DP-24 DataPorts (Fonken, Kitsmiller, Smale, & Nelson, 2012; Watkins et al., 1995). Surgeries were conducted between Zeitgeber time (ZT) 2 and ZT11—with ZT0 being “lights on” and ZT12 “lights off”—and included a T8 laminectomy for all animals. Whereas sham animals received no further surgical intervention, SCI animals received a moderate contusion SCI (150 kdyn, 1-second dwell; Infinite Horizon Impactor) (Gaudet et al., 2017, 2019; Gaudet, Sweet, Polinski, Guan, & Popovich, 2015).

Due to the limited number of telemetry receivers and to minimize potential confounds from conducting experiments with both sexes in overlapping space and time, male and female experiments were performed separately. Male rats (sham, n = 6; SCI, n = 5) were tested first. Note, one transmitter failed to record. Post-surgery care included twice-daily handling (bladder emptying or sham handling) at ZT2 and ZT10, with subcutaneous antibiotic (gentamicin sulfate) and rehydrating saline injections for five days at ZT2. After the male study, the experimental design was improved for the female study. Female rats (sham, n = 6; SCI, n = 6) underwent presurgical handling twice daily to acclimate to procedures. Post-surgery care followed the same protocol as males, except antibiotic/saline injections were given in the afternoon (ZT10) to limit handling-induced disruption of early inactive phase rhythms. Handling was discontinued once animals voided independently, with daily checks and weekly cage changes continuing.

### 2.2. Data acquisition and statistical analysis

All analyses were conducted in *R* (v4.3.2) using *RStudio* (v2024.12.0) (R Core Team, 2022; RStudio Team, 2020). Data wrangling and plotting were performed with *tidyverse* libraries (v2.0.0), particularly *dplyr*, *lubridate*, and *ggplot2*, while figures were assembled using *cowplot* (v1.1.3) (Wickham et al., 2019; Wilke, 2024). Heat map clustering was performed with the *hclust* function from *stats*.

Core temperature values were averaged into 30-minute bins for each subject prior to analysis. Baseline thresholds were calculated as the daytime median temperature plus the interquartile range (IQR). ΔT values were computed as recorded temperature minus the baseline threshold, with negative values set to zero.

Temperature peaks were defined as the highest ΔT within 90 minutes of room entry based on entry/exit logs. Analyses of these values used only the first room entry/stress event of the day.

To test for effects on ΔT values (**Table S1,S2**), a linear mixed-effects model was constructed using *lmerTest* (v3.1-3) with fixed effects for surgery, days post-operative (dpo), sex, and their interactions, and time of day as a covariate (Bates, Mächler, Bolker, & Walker, 2015; Kuznetsova, Brockhoff, & Christensen, 2017). Random intercepts accounted for repeated measures within individual rats. Fixed effects and interactions were evaluated with Type III analysis of variance using Satterthwaite’s method for degrees of freedom via the *anova* function. *Post-hoc* comparisons were performed using estimated marginal means via *emmeans* (v1.10.6) with false-discovery rate (FDR) adjustment for multiple testing (Benjamini & Hochberg, 1995; Lenth, 2024).

To analyze recovery of temperature peaks over time, a linear mixed-effects model was constructed with fixed effects for surgery, dpo (modeled as a natural spline with 2 degrees of freedom), sex, and their interactions. Natural splines (*ns* function from *splines*) were used to capture the asymptotic behavior of temperature recovery (Perperoglou, Sauerbrei, Abrahamowicz, & Schmid, 2019) (**Table S3**). As with the ΔT values, only the first stress interaction of the day was analyzed. Covariates included time of day and daily threshold temperature, with random intercepts for individual rats. Estimated marginal means were used to calculate effect sizes at each dpo, with pairwise comparisons estimating differences between surgery groups while accounting for the model structure (**Table S4**).

## 3. Results

### 3.1. Stress-induced hyperthermia is suppressed by thoracic spinal cord injury

Core temperature is maintained by the hypothalamus and is increased after acute stress via the autonomic nervous system (Nakamura & Morrison, 2022). To establish whether SCI disrupts stress-induced hyperthermia, we analyzed core temperature data measured in male and female Sprague Dawley rats for several weeks before and after a moderate SCI (or sham surgery) (**Fig. 1A-D**). Pre-surgery core temperature during the light phase (ZT0-12) averaged 37.4 ± 0.03°C in males and 37.5 ± 0.02°C in females, while during the dark phase, it averaged 38.0 ± 0.03°C in males and 38.1 ± 0.03°C in females. Stress responses in rats were elicited by procedures required for the study, including post-operative care and other researcher-subject interactions (e.g., cage changes and visual inspection). Prior to surgery, these stress-inducing procedures caused pronounced temperature spikes within ∼30 minutes of room entry (**Fig. 1B**). Females were acclimated to handling pre-surgery; handling elicited an average temperature of 38.4 ± 0.04°C, an increase of 0.9°C above the average female daytime temperature. Similar effects have been reported across species in response to various stimuli (see **Discussion**). These responses are collectively known as stress-induced hyperthermia, and we will refer to them as such throughout this paper (Bouwknecht, Olivier, & Paylor, 2007; Jasim, 2021).

**Figure 1.**
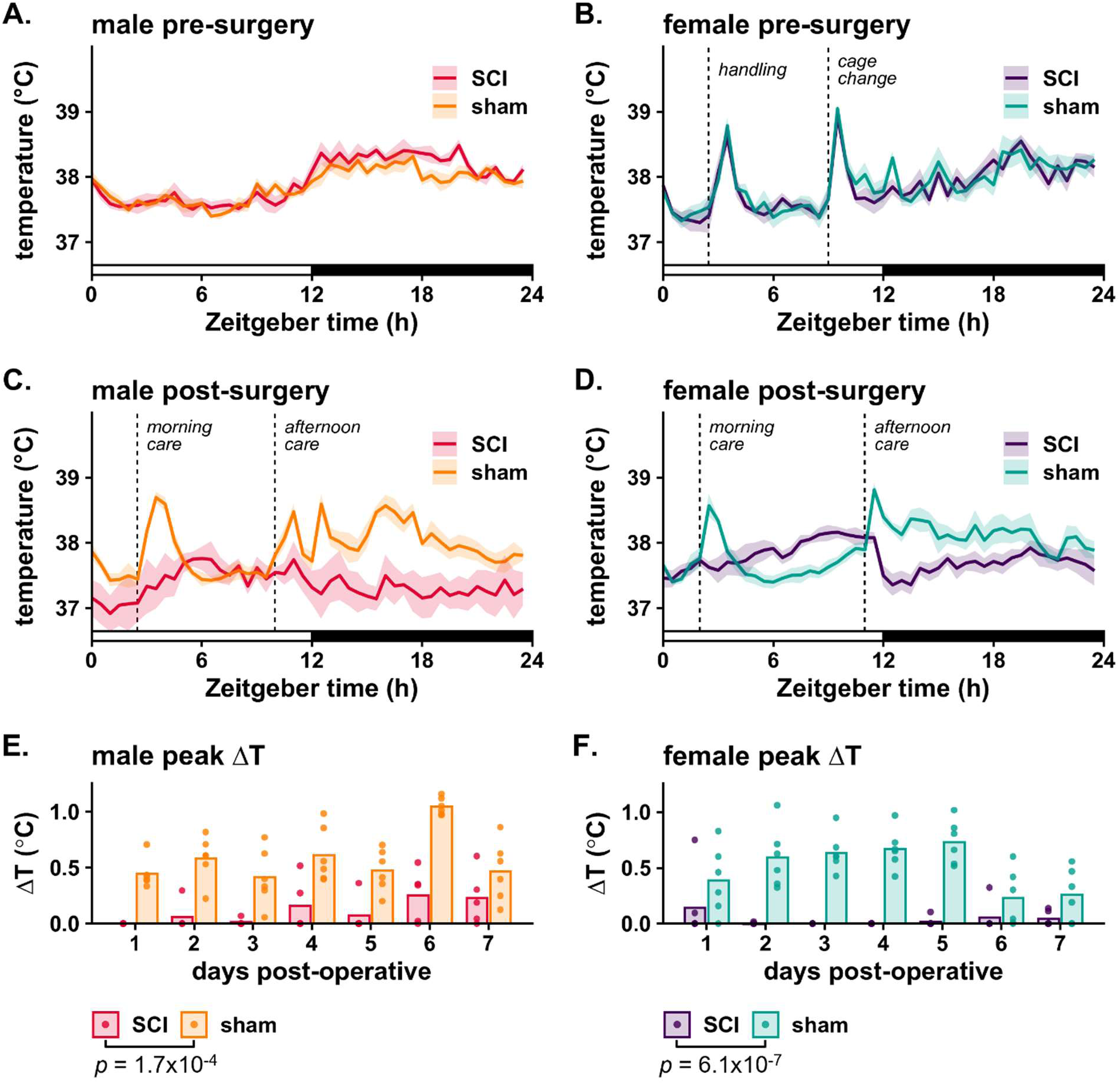
The stress-induced hyperthermic response is abolished acutely after SCI. (**A,B**) Average core temperature (± SEM) for male (**A**) and female (**B**) rats is shown at 1-2 weeks pre-surgery. The x-axis represents Zeitgeber Time (ZT), with ZT0 denoting lights on and ZT12 lights off. Handling/care events are annotated with vertical dashed lines. (**C,D**) Sham and SCI temperatures at 2 dpo – whereas sham rats show typical stress-induced hyperthermia, SCI rats lose this hyperthermic response. (**E,F**) Temperature peaks above background (ΔT) for sham and SCI rats for 7 dpo, with bars representing group means and points marking individual values. SCI disrupts stress-induced hyperthermia across the acute post-injury period. P-values, indicating significant main effects of surgery, were determined via mixed-effects model.

SCI transiently abolished the hyperthermic response (**Fig. 1C,D**). Male and female rats received twice-per-day care during the acute post-surgical phase. Rats that received sham surgery maintained typical hyperthermic responses at these early timepoints (e.g., 2 dpo, **Fig. 1C**). In contrast, SCI rats showed no significant hyperthermia (**Fig. 1D**). For example, over the first three days post-surgery, SCI males reached an average peak temperature of 37.2 ± 0.35°C following care; only 0.3°C above the daily median of 36.9°C. SCI females showed a similar pattern, with an average peak of 37.5 ± 0.32°C and a median of 37.2°C. In contrast, sham-operated rats consistently exhibited stress-induced hyperthermia. Sham males had an average post-care peak of 38.9 ± 0.07°C—0.9°C above the 37.9°C median, while sham females peaked at 38.5 ± 0.01°C, with a median of 37.8°C.

### 3.2. Stress-induced hyperthermia is robustly reduced in the first week after SCI

To investigate the loss of stress-induced hyperthermia acutely after SCI, we calculated the peak temperature change (ΔT) following each morning care event relative to baseline for each animal. This allowed us to isolate the magnitude of the stress-response independently of the previously described differences in daily core temperature and diurnal temperature rhythms (Gaudet et al., 2018). Note, for consistency, in this and subsequent analyses, we limited our observations to the first interaction of the day.

In the acute post-injury phase (1-7 dpo), stress-induced hyperthermia was markedly reduced in SCI animals compared to shams, with minimal ΔT observed following morning care events (**Fig. 1E,F**). Over this period, SCI males showed an average ΔT of 0.11 ± 0.05°C compared to 0.58 ± 0.03°C in sham males, while SCI females had an average ΔT of 0.04 ± 0.02°C compared to 0.5 ± 0.4°C in sham females. Mixed-effects modeling (**Table S1,S2**) revealed a significant main effect of surgery (males: F1, 65.5 = 15.9, p = 1.7×10^-4^; females: F1, 80 = 29.4, p = 6.1×10^-7^), indicating consistently lower ΔT in SCI animals compared to shams throughout the first 7 days post-injury. The lack of a significant surgery × dpo interaction (males: F1, 64 = 0.009, p = 0.93; females: F1, 80 = 1.6, p = 0.21) suggests that SCI animals showed minimal improvement in ΔT across the acute post-injury phase. Day-to-day variation in ΔT was notable even among sham animals, likely driven by differences in care timing and researcher identity and experience level. Thus, incomplete thoracic SCI largely abolished stress-induced hyperthermia throughout the acute post-injury phase.

### 3.3. The stress-induced hyperthermic response gradually recovers over time after incomplete thoracic SCI

To visualize the acute-to-chronic SCI dynamics of stress-induced hyperthermia, we mapped ΔT responses of each rat across seven weeks post-surgery (**Fig. 2A,B**). This approach highlights SCI effects, as well as within– and between-animal variability and distinct day-to-day fluctuations. In rats with sham surgery, the highest ΔT responses occurred due to cage changes (“v” in figure) combined with post-operative care (horizontal lines), followed by care alone, and finally quick visual inspections, which began around 28 dpo once all rats had recovered bladder function. In contrast to sham rats, those with SCI showed dampened stress-induced hyperthermia for up to ∼28 dpo.

**Figure 2.**
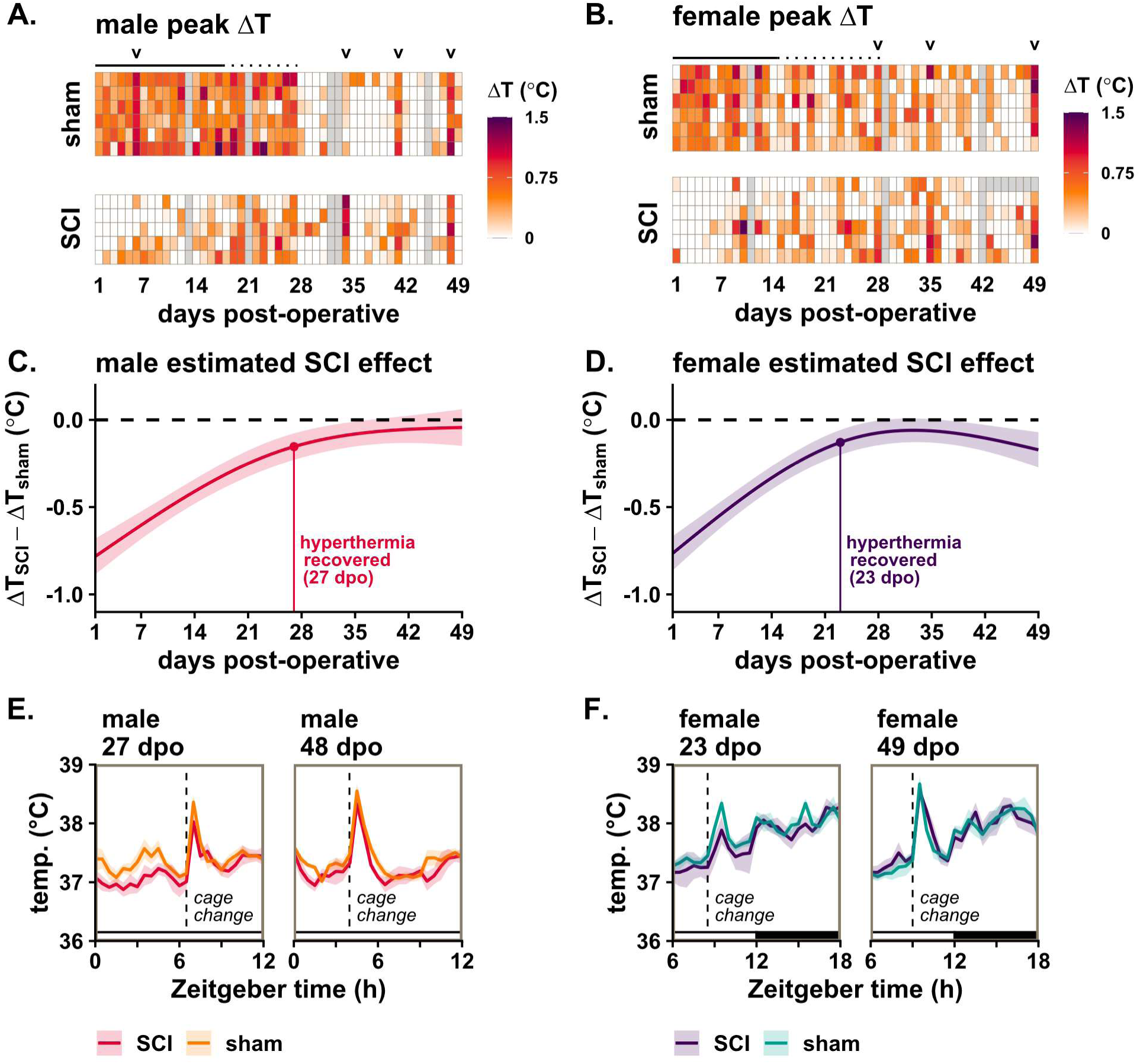
Stress-induced hyperthermia is abolished over acute-to-subacute times after incomplete SCI, and is regained by chronic post-SCI times. (**A,B**) Heatmaps display ΔT (temperature peaks above background) for individual male (**A**) and female (**B**) rats across 7 weeks post-surgery. The first care or interaction event of the day is shown. Cage changes are marked with “v”. Solid lines indicate twice-daily care (1–17 dpo for males and 1–14 for females), while dotted lines mark once-daily care (27 dpo for males and 28 dpo for females). Days with no care events or missing data are shaded in grey. (**C,D**) Modeled recovery curves for the stress-induced hyperthermic response in male and female animals are based on daily estimated marginal means derived from a mixed-effects model. Lines represent the modeled estimates ± standard error. Vertical lines indicate the estimated stress-hyperthermic recovery dates, defined as the first day when SCI stress-induced hyperthermia responses were statistically indistinguishable from sham responses (male SCI vs. male sham: 27 dpo; female SCI vs. female sham: 23 dpo) (*p* > 0.05). (**E,F**) Average core temperatures (± SEM) are shown for sham/SCI males and females on the estimated stress-induced hyperthermia recovery date (left panel, **E** and **F**) and during the final cage change at 7 weeks post-surgery (right panel). Room entries for cage changes are annotated with vertical dashed lines.

At chronic times after incomplete SCI, male and female rats showed more typical stress-induced hyperthermic responses – e.g., SCI rats had robust hyperthermic responses to cage changes around 34-35 and 48-49 dpo (**Fig. 2A,B**).

### 3.4. Stress-induced hyperthermia is restored in rats with chronic incomplete SCI

Next, we sought to capture and quantify the longitudinal effect of SCI on stress-induced hyperthermia. Accordingly, we modeled temperature spike amplitude over the course of the study using a mixed-effects approach, including surgery, dpo, and sex as factors, with time-of-day and baseline daily temperature as covariates. As established from the ΔT data, there was a strong interaction between surgery and dpo (F2, 997.7 = 48.20, p < 2.2×10^-16^), reflecting the loss of the stress-induced hyperthermic response immediately after SCI and its gradual recovery over time. Despite differences in acclimation and care timing, sex showed little effect in our model, with no significant main effect (F1, 120.3 = 0.09, p = 0.759), no significant interaction with surgery (F1, 117.4 = 0.02, p = 0.900), and no significant effect of the surgery × dpo interaction (F2, 997.1 = 0.92, p = 0.399; **Table S3**).

Finally, to estimate the recovery timing of the stress-induced hyperthermic response, we analyzed the modeled daily mean effect of SCI on the hyperthermic response in males and females (**Fig. 2C,D; Table S4**). These effects were derived from a *post-hoc* analysis of estimated marginal means. Date of recovery was estimated as the first day when the modeled temperature responses of SCI animals were statistically indistinguishable from those of sham animals (p > 0.05). By this metric, SCI males had largely recovered stress-induced hyperthermia by 27 dpo and SCI females by 23 dpo, which is consistent with the broader model results showing time-dependent surgery effects with minimal evidence for sex-specific differences. Raw temperature traces at these time points illustrate the reemergence of stress-induced hyperthermia, though attenuated compared to shams (at 27 dpo (males) and 23 dpo (females); **Fig. 2E,F** left panels). By 7 weeks, the response appears fully restored (at 48 dpo (males) and 49 dpo (females); **Fig. 2E,F** right panels). Overall, these results reveal that incomplete thoracic SCI abolishes stress-induced hyperthermia in male and female rats acutely after injury, and that the hyperthermic response regains typical amplitude chronically after SCI.

## 4. Discussion

Stress-induced hyperthermia is a physiological response observed in various animal species, characterized by a transient increase in core body temperature following exposure to acute stressors (Vinkers et al., 2010). In laboratory mice and rats, stressors such as handling/restraint, exposure to novel objects or environments, and social defeat can lead to a significant rise in core body temperature, typically ranging from 0.5°C to 1.5°C (Borsini, Lecci, Volterra, & Meli, 1989; Bouwknecht, Olivier, & Paylor, 2007; Lkhagvasuren, Nakamura, Oka, Sudo, & Nakamura, 2011; Vinkers, van Oorschot, Olivier, & Groenink, 2009). Similarly, in humans, acute psychological stress can trigger transient increases in core body temperature (i.e., “psychogenic fever”), whereas chronic stress can lead to persistent low-grade hyperthermia (∼38°C) (Oka, 2015; Vinkers, Groenink, et al., 2009). Unlike conventional fever, stress-induced hyperthermia is driven by autonomic activation rather than inflammatory signals, as evidenced by its resistance to antipyretics and attenuation by sympathetic blockers and anxiolytics (Bouwknecht et al., 2007; Oka, Oka, & Hori, 2001; Olivier et al., 2003; Vinkers, Groenink, et al., 2009; Watkins et al., 1995).

In this study, we continuously monitored core temperature in rats and observed similar stress-induced hyperthermic responses following handling, cage changes, and other necessary pre– and post-surgery interventions (**Fig. 1A-D**). In the first week after surgery, the hyperthermic response was largely abolished in SCI rats while remaining unaltered in sham rats (**Fig. 1E,F**). Thermoregulatory dysfunction is a well-documented consequence of SCI in humans, often leading to challenges in maintaining core body temperature (Downey, Chiodi, & Darling, 1967; Guttmann, Silver, & Wyndham, 1958; Price & Campbell, 2003). Broader autonomic disruptions, such as autonomic dysreflexia—a potentially life-threatening condition—are also commonly associated with SCI (Fauss, Hudson, & Grau, 2022; Furlan & Fehlings, 2008; Partida, Mironets, Hou, & Tom, 2016). Similar thermoregulatory and autonomic disruptions occur in rodent models of SCI, including our own observations of altered circadian temperature rhythms and general hypothermia during the first 1-3 dpo (Chiu & Lee, 2021; Gaudet et al., 2018; Kalincik, Jozefcikova, Waite, & Carrive, 2009; Laird, Carrive, & Waite, 2006; Trueblood, Iredia, Collyer, Tom, & Hou, 2019). To our knowledge, however, this is the first report detailing that SCI suppresses stress-induced hyperthermic responses.

Thermoregulation is broadly controlled by the hypothalamus, which integrates diverse physiological and environmental signals to maintain core body temperature. In the context of stress-induced hyperthermia, the hypothalamus processes acute stress signals from higher brain centers and coordinates autonomic outputs to peripheral effectors including brown adipose tissue and cutaneous vasculature. This response is further amplified by the adrenal glands, which release stress hormones to enhance thermogenic and cardiovascular effects (Lkhagvasuren et al., 2011; Nakamura & Morrison, 2022; Ootsuka, Blessing, & Nalivaiko, 2008; Wulf & Tom, 2023). These pathways rely on descending projections through the medullary raphe and spinal cord to regulate activation of the sympathetic nervous system. The T8 contusion SCI we used in this study likely disrupts these descending pathways, impairing the autonomic signals required for the hyperthermic response. Specifically, damage to the intermediolateral (IML) cell column at this level could block thermogenic activation of brown adipose tissue or hinder vasoconstriction, resulting in the observed loss of hyperthermic responses in SCI animals (Wulf & Tom, 2023). Additionally, given that stress-induced hyperthermia is partially mediated via vagal afferents (Watkins et al., 1995), SCI-related autonomic disruptions may impair this pathway, altering how peripheral stress and inflammatory signals reach central thermoregulatory circuits. Compounding this, the systemic inflammation triggered by SCI could further dysregulate vagally mediated immune-brain communication, exacerbating impairments in stress-induced thermoregulation (DiSabato et al., 2024).

SCI-elicited loss of stress-induced hyperthermia at 1-7 dpo (**Fig. 1E,F**) is likely driven by various acute phase pathological processes acting in concert to disrupt essential autonomic pathways. Immediately following SCI, excessive release of neurotransmitters from damaged neurons overstimulates receptors, causing ionic imbalances, oxidative stress, and further neuronal damage/death (termed excitotoxicity; (Ahuja et al., 2017; Kwon, Tetzlaff, Grauer, Beiner, & Vaccaro, 2004; Schanne, Kane, Young, & Farber, 1979). This is accompanied by a robust neuroinflammatory response, which peaks within 3-7 dpo. Activation of resident microglia and infiltration of peripheral immune cells lead to the release of pro-inflammatory cytokines and reactive oxygen species, further exacerbating tissue damage and impairing neuronal function (Gaudet & Fonken, 2018; Hellenbrand et al., 2021). Together, these processes likely contribute to disruption of neural circuits critical for autonomic regulation during the acute phase.

Although neuroinflammation can persist for months to years following SCI, its attenuation after the acute phase, coupled with the clearance of excitotoxic factors, may allow spared neurons to regain normal function. These changes, occurring within weeks post-injury, likely support the recovery of autonomic pathways and contribute to the gradual restoration of stress-induced hyperthermia observed in our study (**Fig. 2**).

In contrast to the post-SCI recovery of the stress-induced hyperthermic response (**Fig. 2C–F**), locomotor function never fully recovers in our contusion SCI model (Gaudet et al., 2017, 2018). Rats assessed in parallel to this study, plateaued at a score of ∼10 on the Basso, Beattie, and Bresnahan (BBB) scale—indicating occasional weight-supported plantar steps without coordinated forelimb-hindlimb movements—from 2 to 6 weeks post-injury (Basso, Beattie, & Bresnahan, 1995; Gaudet et al., 2018; Ma, Basso, Walters, Stokes, & Jakeman, 2001). Notably, the stabilization of motor function occurs on a similar timeline as the recovery of stress-induced hyperthermia, suggesting overlapping repair processes. The differences in recovery outcomes likely reflect the distinct anatomical locations of the descending motor and autonomic pathways. In rodents, the corticospinal tract, which is the primary motor pathway, is housed within the dorsal funiculus (Watson & Harrison, 2012), a region that sustains extensive damage in our T8 contusion model. In contrast, the IML is positioned between the dorsal and lateral horns, further away from the primary impact site. This localization may afford it a degree of protection and facilitate a greater degree of functional recovery. Furthermore, since the IML spans the T1 to L2 spinal segments (Wulf & Tom, 2023), much of the sympathetic outflow remains intact above the T8 injury level in our model. This preserved circuitry may provide a structural basis for the more complete recovery of stress-induced hyperthermia, allowing spared pathways to compensate for lost function below the lesion.

Our findings demonstrate that SCI severely disrupts the stress-induced hyperthermic response during the acute post-injury phase, with gradual recovery to pre-injury levels by the study’s conclusion (7 weeks post-SCI). The primary limitations of this study stem from its post-hoc, observational nature, as the original experiments focused on circadian temperature rhythms rather than SCI’s effects on stress-induced hyperthermia. Consequently, uncontrolled factors—including stressor type (e.g., researcher-elicited bladder voiding, subcutaneous injections, observation, cage change), as well as the timing, frequency, and duration of care events—likely influenced the observed responses. Furthermore, due to the limited number of telemetry systems, males and females were assessed in separate cohorts using slightly altered methodology, limiting our ability to more-thoroughly assess the effects of sex. Future studies with precisely controlled stress-induced hyperthermia methodologies may help clarify subtle sex differences, while experiments targeting specific pathways could identify the circuits most vulnerable to SCI and their role in autonomic recovery.

Given that incomplete T8 SCI transiently abolishes stress-induced hyperthermia, future studies should explore how SCI of different severities (e.g., complete SCI) or at different levels (e.g., T3 vs. T8) affect the extent and duration of autonomic dysfunction. Indeed, previous studies have revealed that complete high-thoracic SCI more robustly disrupts autonomic function, contributing to conditions such as autonomic dysreflexia and immune system disruption (Mironets et al., 2018; Noble et al., 2022; Rodgers, Kigerl, Schwab, & Popovich, 2022; Zhang et al., 2013). These findings suggest that higher-level injuries could have wide-ranging implications for the regulation of stress-induced hyperthermia and other autonomic functions, potentially mediated by both neural and immune mechanisms.

## 5. Conclusions

We explored the stress-induced hyperthermia response in rats prior to and after SCI. SCI in female and male rats abolished stress-induced hyperthermia. This SCI-driven autonomic dysfunction was particularly striking acutely after SCI; stress-induced hyperthermia gradually re-emerged in the chronic phase (by 3-7 weeks post-SCI).

Interestingly, reinstatement of these autonomic responses contrasts with permanent locomotor impairments in this same rat SCI model, likely due to distinct pathways and plasticity between the two systems. Overall, our data reveal that incomplete thoracic SCI severely impairs a typical thermoregulatory response, stress-induced hyperthermia. These results underscore the importance of studying and treating various autonomic outcomes after SCI. Future studies should reveal mechanisms underlying autonomic dysfunction and plasticity after SCI.

## Acknowledgements

The authors thank the Muenzinger and Wilderness Place husbandry staff for excellent animal care at CU-Boulder. At the University of Texas at Austin, the authors thank the staff at the Animal Resource Center.

## Abbreviations

SCI: Spinal cord injury
ANS: Autonomic nervous system
ZT: Zeitgeber time
IQR: Interquartile range
ΔT: Change in temperature above baseline
IML: Intermediolateral Cell Column
BBB: Basso, Beattie, and Bresnahan Score
FDR: False Discovery Rate
LMM: Linear Mixed-effects Model

